# ATOMDANCE: kernel-based denoising and allosteric resonance analysis for functional and evolutionary comparisons of protein dynamics

**DOI:** 10.1101/2023.04.20.537698

**Authors:** Gregory A. Babbitt, Madhusudan Rajendran, Miranda L. Lynch, Richmond Asare-Bediako, Leora T. Mouli, Cameron J. Ryan, Harsh Srivastava, Patrick Rynkiewicz, Kavya Phadke, Makayla L. Reed, Nadia Moore, Maureen C. Ferran, Ernest P. Fokoue

**Affiliations:** Thomas H. Gosnell School of Life Sciences, Rochester Institute of Technology, Rochester, NY 14623, USA; Hauptmann Woodward Medical Research Institute, Buffalo, NY, USA; McQuaid Jesuit High School Computer Club, Rochester, NY 14618, USA; New York University, Rochester, NY 14618, USA; School of Mathematical Sciences, Rochester Institute of Technology, Rochester, NY 14623, USA

**Keywords:** comparative method, machine learning, molecular dynamics simulation, molecular evolution, resonance, thermal noise, allosteric regulation

## Abstract

Comparative methods in molecular biology and molecular evolution rely exclusively upon the analysis of DNA sequence and protein structure, both static forms of information. However, it is widely accepted that protein function results from changes in dynamic machine-like motions induced by molecular interactions, a type of data for which comparative methods of analysis are challenged by the large fraction of protein motion created by random thermal noise in the surrounding solvent. Here, we introduce ATOMDANCE, a suite of statistical and kernel-based machine learning tools designed for comparing and denoising functional states of protein motion captured in time-series from molecular dynamics simulations. ATOMDANCE employs interpretable Gaussian kernel functions to compute site-wise maximum mean discrepancy (MMD) between learned features of motion representing two functional, malfunctional or evolutionary protein states (e.g. bound vs. unbound, wild-type vs. mutant). ATOMDANCE derives empirical p-values identifying functional similarity/difference in dynamics at each amino acid site on the protein. ATOMDANCE also employs MMD to contextually analyze potential random amino-acid replacements thus allowing for a site-wise test of neutral vs. non-neutral evolution in the divergence of dynamic function in protein homologs. Lastly, ATOMDANCE also employs mixed-model ANOVA combined with graph network community detection to identify functional shifts in protein regions that exhibit time-coordinated dynamics or resonance of motion across sites. Here, we demonstrate the utility of the software for identifying key sites involved in dynamic responses during functional binding interactions involving DNA, small molecule drugs, and virus-host recognition. We also demonstrate its utility in understanding dynamic resonance changes occurring during the allosteric activation of a pathogenic protease. ATOMDANCE offers a user-friendly interface and only requires an input structure, topology and trajectory files for each of the two proteins being compared (i.e .pdb, .prmtop, and .nc). A separate interface for generating molecular dynamics simulations via open-source tools is also offered.

Lay Audience Summary – ATOMDANCE is a suite of software pipelines controlled by a single user interface and designed to comprehensively simulate, calculate and compare protein motions between two functional or evolutionary states while controlling for random noise. It is useful for finding amino acid sites on a given protein that are important in binding other proteins, DNA, or drugs/toxins. It can also be used to assess the effect of genetic mutation on protein motion, and identifies regions of sites on proteins that tend to move together or ‘resonate’ as a whole unit or community.

## Introduction

Protein sequences are typically less conserved over evolutionary timescales than are protein structures (Illergård et al., 2009; Sousounis et al., 2012). This implies that the most important functional information contained in protein sequences is often manifested at the level of protein structure and the motion or dynamics of their structural interactions. However, while protein specificity during molecular interactions has often been compared to a static lock and key (Fischer, 1894), proteins and many of the binding partners they interact with exhibit extensive soft matter physical properties, implying that protein-ligand function cannot be entirely understood through the analysis of protein-ligand structure alone (Koshland, 1958; Tripathi and Bankaitis, 2017). Often proteins are described as analogous to nanoscale-sized machines (Abendroth et al., 2015; Flechsig and Mikhailov, 2019; Strong, 2004), where non-random repetitive motions, which should be discernible from random thermal noise, are key characteristics of protein function. Thus the functional evolution of the cell must depend heavily upon how protein structures alter or shift their dynamic motions during important motor functions and logic gating functions required when proteins and other biological macromolecules collectively assemble to form key metabolic or regulatory pathways (Babbitt et al., 2016).

Comparative methods of analysis are well developed for protein sequences and structures in the disciplines of phylogenetics (Cornwell and Nakagawa, 2017), molecular evolution (Suzuki, 2010) and structural biology (Kufareva and Abagyan, 2012). And they are deeply rooted in experimental biology and its association with the historical development of the field of modern statistics (Parolini, 2015). In molecular evolution and comparative genomics, it is important to note that these methods are often applied in a specific site-wise manner as evolutionary changes due to genetic mutation tend to act independently at individual sites over time. Site-wise analyses of root mean square deviation (RMSD) of superimposed protein chains are also very common in structural biology. However, site-wise comparative methods for application to molecular dynamics are only now beginning to be developed (Babbitt et al., 2020, 2018). Important types of site-wise comparisons derived from two different molecular dynamics (MD) simulations might include (A) comparing the same amino acid site in two functional states (e.g. bound vs unbound to drugs, toxins, nucleic acids, or other proteins), (B) comparing the same sites at two different temperatures in an investigation of thermostability, (C) comparing the same amino acid site in two different evolutionary lineages or before and after a significant mutation event, or (D) comparing the same amino acid site in two different epigenetic states (e.g. involving phosphorylation or methylation). Alternatively, important types of site-wise comparisons taken from the same MD simulation could include comparing the dynamics of two adjacent or non-adjacent amino acid sites over time in order to ascertain how coordinated or ‘resonant’ they are in their dynamic behavior, possibly indicating native contact of nearby sites or even allosteric regulation involving more distant sites. Being able to make site-wise determinations of similarities and differences in protein dynamics has potential application to the fields of computational pharmacology and vaccine development as they interface with the basic science of molecular evolution. Potential applications include the problems of identifying/predicting single protein sites involved in the evolution of vaccine (Rajendran et al., 2022) or drug escape (Rajendran et al., 2023) and/or the problem of whether a generic drug will interact similarly enough to a patented product across humans and model lab animals given their respective genetic and epigenetic differences in protein targets.

A major challenge with attempting to functionally analyze the dynamic trajectories of atoms in MD simulations is caused by the large fraction of motion in the system that is simply due to random thermal noise. This noise can obscure the non-random machine-like motions most connected to protein function. This is especially problematic in explicit solvent based MD approaches, which more accurately replicate protein dynamics than implicit solvent methods (Darden et al., 1993; Petersen, 1995; Wang et al., 2012). Thermal noise in explicit solvent MD simulation is caused by the random collision of water molecules and free ions with the protein chains. Past methods of comparative molecular dynamics analysis have relied upon a large amount of resampling of the atom trajectories to be able to resolve site-wise functional differences in dynamics caused by binding interactions and mutations (Babbitt et al., 2020, 2018). However, because a machine learning algorithm cannot be expected to learn from noise, it could theoretically provide a useful method of detecting signal from noise (i.e. denoising) when analyzing functional differences in protein states captured in MD simulations. Despite their promise for potentially filtering random from non-random motion in MD simulation, site-wise machine learning approaches remain largely unexplored for this purpose; however see (Babbitt et al., 2020) for one approach to identifying regions of conserved dynamics.

The motion of proteins over relatively short time scales in MD simulations is largely harmonic and therefore we propose that they can be best addressed with a Gaussian kernel-based approach to machine learning. Because proteins are polymers, the motions of constituent atoms are often restricted by steric constraints caused by the protein folding. This creates unequal variances in the degree of atom fluctuation (i.e. directionless magnitude of motion) across sites. This harmonic oscillation with unequal variance in atom motion in protein polymer chains implies that a machine learning approach that incorporates a Gaussian kernel function might perform well when tasked with identifying functional or evolutionary differences when comparing noisy explicit solvent MD simulations. A Gaussian process kernel learner also has an advantage in that it is much more interpretable than black box methods such as support vector machine and neural networks when applied to many physical systems (Ponte and Melko, 2017). This interpretability might be very important to biomedical researchers when navigating a rapidly evolving regulatory landscape regarding the application of machine learning to drug discovery, often aiming to decipher complicated protein target interactions with a large variety of small molecule compounds that might outcompete naturally occurring compounds for binding to these targets.

Here we introduce the ATOMDANCE statistical machine learning post-processor for comparative molecular dynamics; a software suite for interpretable machine learning comparison and allosteric resonance analysis of functional protein dynamics, performed at individual site-wise resolution. The ATOMDANCE software suite provides researchers with a user-friendly graphical interfaced computational platform for supplementing comparative sequence/structure analyses with important information about the functional motions of proteins undergoing complex interactions with DNA/RNA, drugs, toxins, natural ligands, or other proteins. It can also address how genetic mutation and/or molecular evolution has altered functional motion as well as how interactions in the motion across sites resonate (i.e. are coordinated over time) and how site resonance might shift during mutation and/or functional interaction with other molecules in the cell. ATOMDANCE interfaces automatically with UCSF ChimeraX to produce color-mapped structural images as well as graph plotting with a python implementation of ggplot (i.e. plotnine module) using a modern PyQt 5 graphical user interface.

## Summarized Methods

See the Supplemental Methods file for more detailed descriptions about these analyses summarized below.

ATOMDANCE offers 4 main analysis pipelines.

(A) DROIDS 5.0 for direct site-wise comparison of protein dynamics

The site-wise average differences and Kullback-Leibler (KL) divergences in atom fluctuation are reported similarly to the DROIDS 4.0 method previously developed by our lab group (Babbitt et al., 2020, 2018). In ATOMDANCE, we offer the same analysis in a completely python-based application as DROIDS 5.0. Sites with significantly different dynamics are identified with a multiple test corrected two sample Komogorov-Smirnov test. While older versions of DROIDS were developed as a perl plus R language pipeline that interfaced directly with licensed Amber software, DROIDS 5.0, like all of ATOMDANCE runs exclusively in python code, and now executes independently of MD simulation software.

(B) maxDemon 4.0 for denoised *<u>functional</u>* comparisons of protein dynamics

The maxDemon 4.0 program in ATOMDANCE is trained on feature vectors of local atom fluctuations derived from the molecular dynamics trajectories of proteins in two functional states (e.g. bound vs. unbound or wild-type vs. mutant). The comparison between MD simulations at given sites are reported as maximum mean discrepancy (MMD) in the reproducing kernel Hilbert space (RKHS). Hypothesis tests for significance of functional dynamic changes reported via MMD are also provided using a bootstrapping approach. A graphical summary of the method is shown in Supplemental Figure 1.

(C) maxDemon 4.0 for denoised *<u>evolutionary</u>* comparisons of protein dynamics

For a test of neutral evolution on protein dynamics, the MMD of amino acid replacements observed on orthologs is compared to a neutral model of the MMD of random pairs of differing amino acids at different sites on the two protein simulations being compared. This allows for the identification of potential sites where natural selection has either functionally conserved or adaptively altered the local molecular dynamics of the protein.

(D) Choreograph 2.0 – classical statistical identification and comparison of dynamics that are coordinated across different sites on a given protein

The ChoreoGraph 2.0 program in ATOMDANCE offers a site-wise mixed-model ANOVA and graph network community detection analysis of time interaction (i.e. resonance) for the identification of regions of amino acid sites with high of coordination in their respective dynamic states. The resonance communities detected in both functional protein states are compared using bootstrapped resampling of graph network connectivity and two methods of defining graph network non-randomness. These are the (A) probability of two sites being in the same resonance community and the (B) distance of the graph network from a random graph defined by the Erdos-Renyi model.

ATOMDANCE is intuitive and user-friendly, providing a simple graphical user interface (GUI) that only requires structure, topology, and trajectory files (.pdb, .prmtop, .nc) for the two molecular dynamics simulations being compared (Figure 1). It is entirely python-based and outside of this it only requires UCSF ChimeraX for molecular visualization and the popular cpptraj library for resampling calculations (Goddard et al., 2018; Pettersen et al., 2021; Roe and Cheatham, 2013). ATOMDANCE is also supplemented with an optional GUI (Supplemental Figure 2) for generating simulations via open-source tools (i.e. openMM and AmberTools) (Case et al., 2005; Eastman et al., 2017). However it can also potentially be used with files generated using NAMD (qwikMD), CHARMM, or the licensed version of Amber (Case et al., 2005; Jo et al., 2008; Phillips et al., 2020; Ribeiro et al., 2016). We provide an optional GUI for generating multiframe PDB file movies using UCSF ChimeraX (Pettersen et al., 2021) where the motions of the protein system are colored and augmented in accordance with the MMD. Examples of these movies can be seen in an introductory video available at https://people.rit.edu/gabsbi/img/videos/MMDmovie.mp4

**Figure 1.**
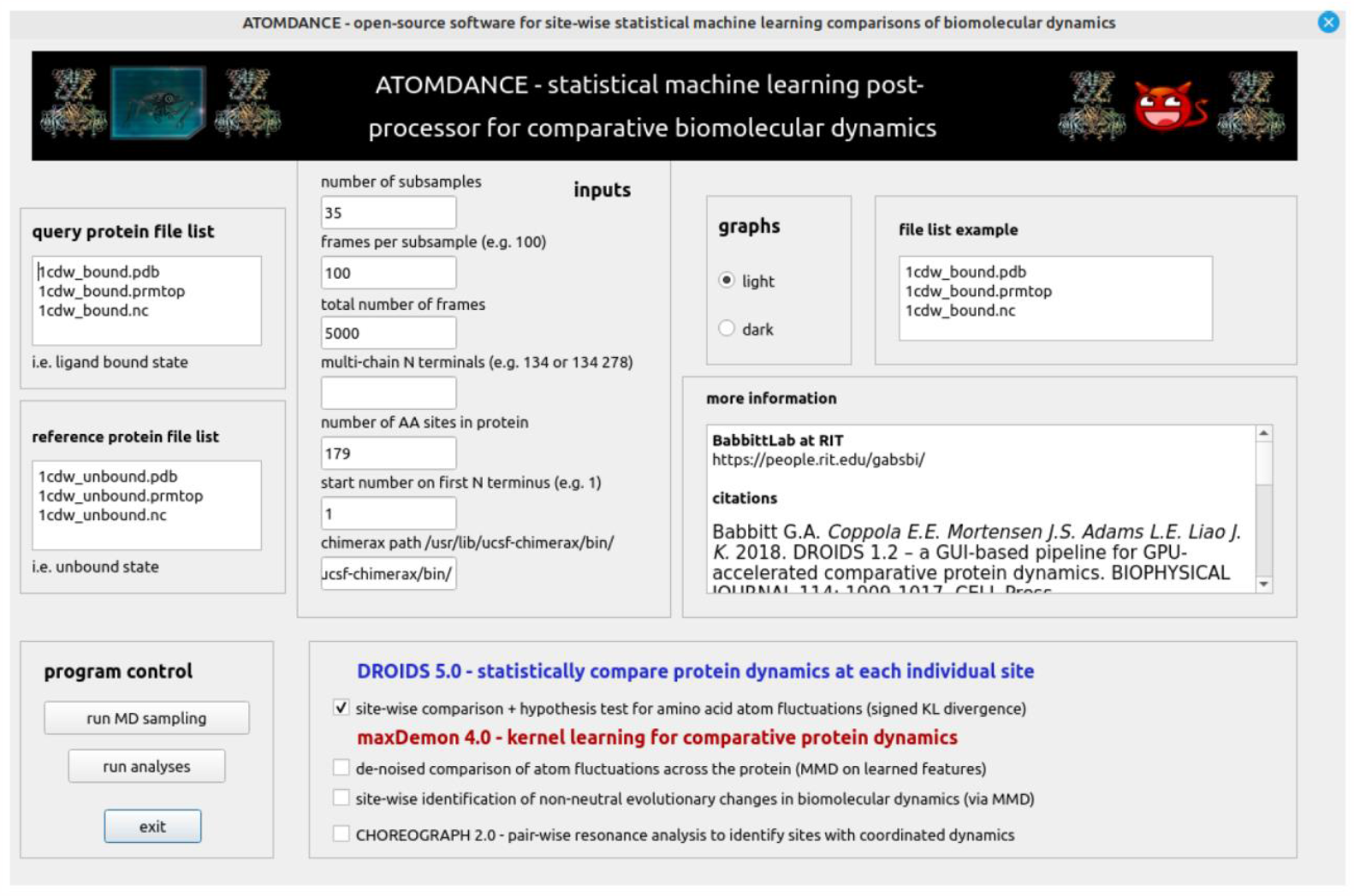
The Graphical User Interface (GUI) for the ATOMDANCE statistical machine learning post-processor for comparative protein dynamics. Each comparison requires six files; two .pdb structure files, two .prmtop topology files, and two .nc trajectory files…each representing one of the two functional or evolutionary states being compared (i.e. bound vs. unbound or before vs. after mutation). The ATOMDANCE GUI for generating molecular dynamics simulations using open-source software is shown in Supplemental Figure 1.

ATOMDANCE is available at GitHub/GitHub pages

https://github.com/gbabbitt/ATOMDANCE-comparative-protein-dynamics

https://gbabbitt.github.io/ATOMDANCE-comparative-protein-dynamics/

and as a docker container here

https://github.com/patrynk/atomdance-docker

Examples presented in this manuscript were generated from structure, topology, and trajectory files deposited here

https://zenodo.org/record/7679282#.Y_wIK9LMJ9A

DOI 10.5281/zenodo.7679282

## Results

To demonstrate the utility of ATOMDANCE, we present a comparison of the unfiltered site-wise divergences in atom fluctuation (Figure 2) to the denoised site-wise discrepancy in learned features of local atom fluctuation (Figure 3), presented in four examples of functional binding interactions. These four examples include (A) DNA-bound vs. unbound TATA binding protein (PDB: 1cdw)(Nikolov et al., 1996), (B) sorafenib-bound vs. unbound B-Raf kinase domain (PDB: 1uwh)(Wan et al., 2004), (C) SARS-CoV-2 viral bound vs. unbound angiotensin-converting enzyme 2 (ACE2) protein (PDB: 6m17)(Yan et al., 2020), and (D) the allosteric activated (i.e. InsP6 bound) vs inactivated (i.e. unbound) Vibrio cholera toxin RTX cysteine protease domain (PDB: 3eeb)(Lupardus et al., 2008). The root mean square fluctuation plots for these comparisons are shown in Supplemental Figure 3.

**Figure 2.**
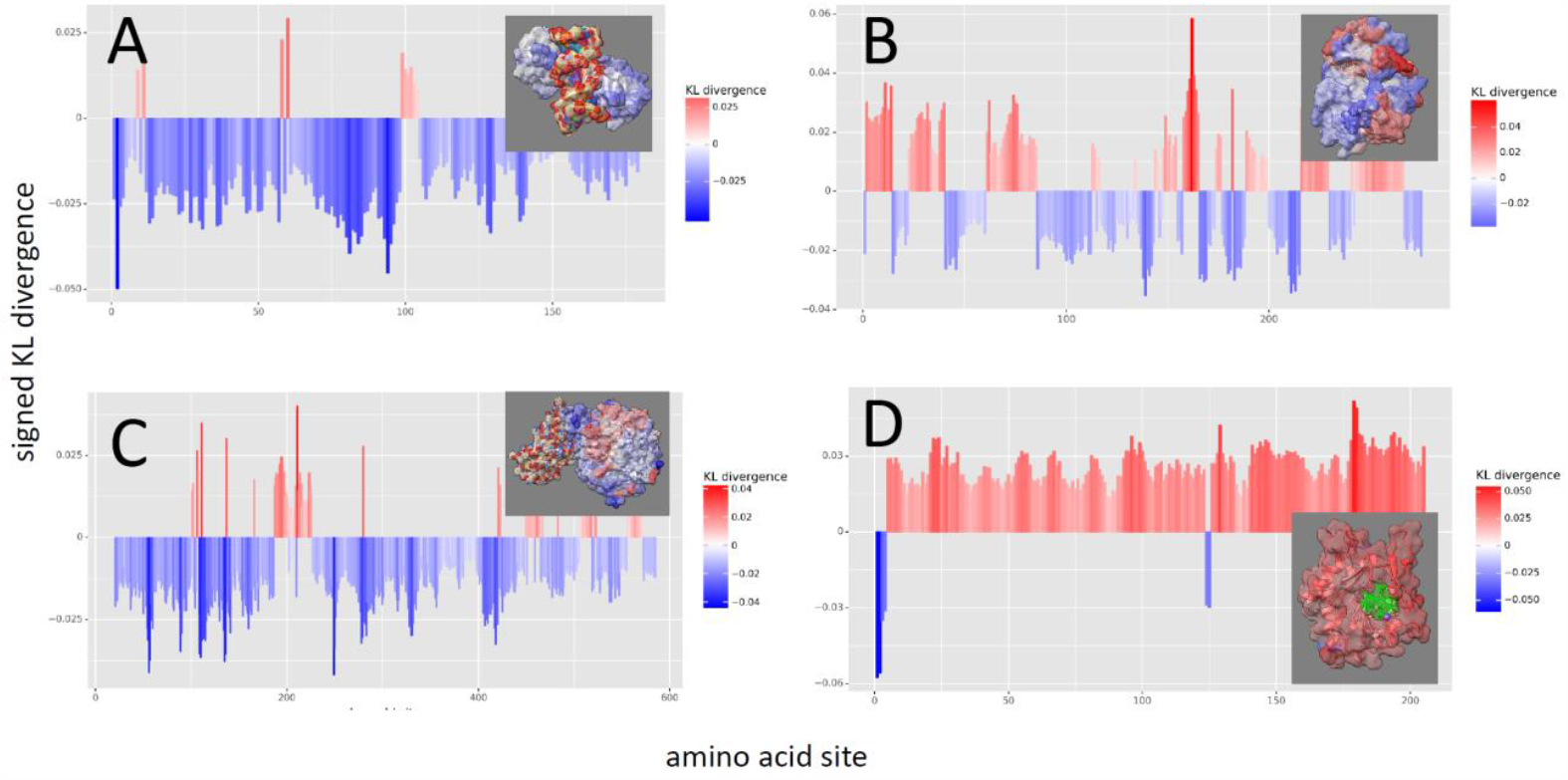
DROIDS 5.0 analysis of site-wise divergence metrics in local atom fluctuation when comparing molecular dynamics simulations of functionally bound vs. unbound target proteins. The comparisons include (A) DNA-bound vs. unbound TATA binding protein (PDB: 1cdw), (B) sorafenib-bound vs. unbound B-Raf kinase domain (PDB: 1uwh), (C) SARS-CoV-2 viral bound vs. unbound angiotensin-converting enzyme 2 (ACE2) protein (PDB: 6m17), and (D) the allosteric activated (i.e. InsP6 bound) vs inactivated (i.e. unbound) Vibrio cholera toxin RTX cysteine protease domain (PDB: 3eeb). Signed symmetric Kullback-Leibler (KL) divergence in atom fluctuation indicates sites where motion is dampened during binding (blue) and where motion is amplified (red). Note that while binding typically dampens atom fluctuation locally or even globally, in the case of this example of allostery (D) it actually amplifies atom fluctuation globally.

**Figure 3.**
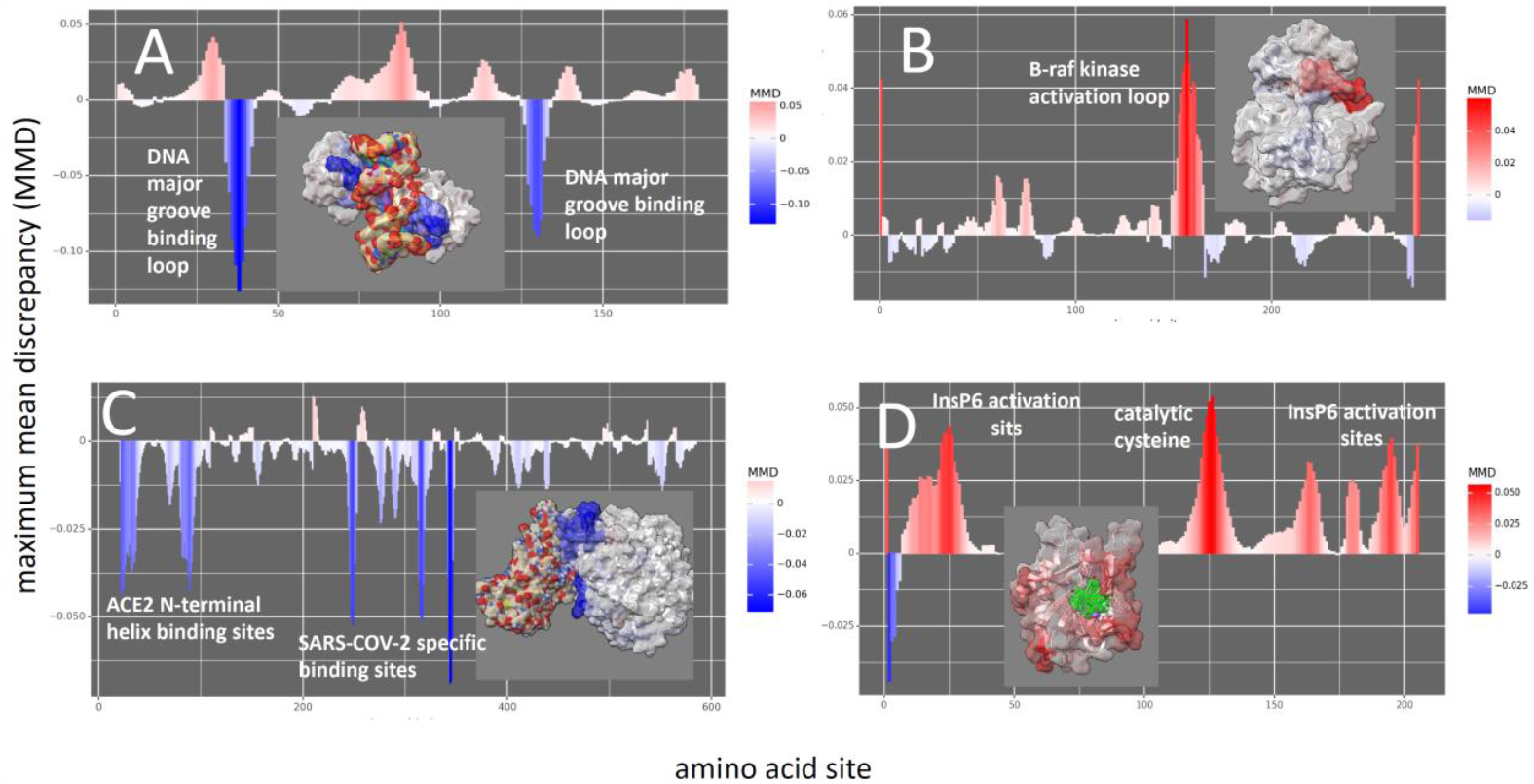
maxDemon 4.0 analysis of signed maximum mean discrepancy (MMD) learned features regarding local atom fluctuation when comparing molecular dynamics simulations of functionally bound vs. unbound target proteins. The comparisons include (A) DNA-bound vs. unbound TATA binding protein (PDB: 1cdw), (B) sorafenib-bound vs. unbound B-Raf kinase domain (PDB: 1uwh), (C) SARS-CoV-2 viral bound vs. unbound angiotensin-converting enzyme 2 (ACE2) protein (PDB: 6m17), and (D) the allosteric activated (i.e. InsP6 bound) vs inactivated (i.e. unbound) Vibrio cholera toxin RTX cysteine protease domain (PDB: 3eeb). Signed MMD in atom fluctuation indicates sites where motion is dampened during binding (blue) and where motion is amplified (red). Note that the kernel-based learning applied to local atom fluctuation (i.e. signed MMD) removes the overall effect of differences in thermal noise present in divergence metrics (Figure 2) from the functional comparison and so much better isolates the binding sites themselves. Close-up views of the color-mapped structures are shown in Supplemental Figure 3.

The first example of comparative protein dynamics analyses conducted with ATOMDANCE investigated the functional effect of DNA binding to TATA binding protein (TBP; PDB 1cdw) by the site-wise comparison of atom fluctuation of TBP in both its DNA bound and unbound state. Figure 2A shows both color-mapped protein surface and site-wise plot of the KL divergence in fluctuation (i.e. DROIDS 5.0). This protein binds quite strongly as is evidenced by a general dampening of fluctuation across the protein (in blue). Supplemental Figure 4 demonstrates alternative plots of the TBP results generated by ATOMDANCE showing site-wise atom fluctuation profiles and average differences in fluctuation colored by amino acid type. The comparison of machine learning derived MMD (i.e. maxDemon 4.0; Figure 3A) clearly captures the key sites of the functional interaction; two loops of the protein that interact directly with the major groove of the DNA (in blue) (Nikolov et al., 1996). Close-up views of all MMD color-mapped structures are given in Supplemental Figure 5 with TBP in panel A. Supplemental Figure 6 shows the TBP MMD plot colored by amino acid type and bootstrapped empirical p-values. The second example of the application of MMD captures the functional amplification of atom fluctuation by the activation loop (shown in red) of BRAF kinase upon the binding of the drug sorafenib in the ATP binding pocket of the kinase domain (PDB 1uwh; Figure 3). In this example the interaction of ATP or ATP-competitive antagonists like the cancer drug sorafenib clearly amplify the motion in the activation loop as can be observed in both the unfiltered and denoised dynamics (Figure 2B and Figure 3B) (Wan et al., 2004). While sorafenib binds the site stronger than ATP, thus interrupting the MAPK pathway triggering cell proliferation in tumorogenesis, this amplification of the activation loop by the drug may be functionally related to the hyperactivation of MAPK in surrounding normal cells, leading to cancer recurrence (Babbitt et al., 2022).

A third example also demonstrates the utility of the DROIDS 5.0 KL divergence and maxDemon 4.0 MMD to investigate the protein-protein interaction between the viral SARS-CoV-2 receptor binding domain (RBD) and its human protein target angiotensin converting enzyme (ACE2)(PDB 6m17) (Yan et al., 2020). While the unfiltered divergence in viral-bound vs. unbound dynamics exhibits general dampening of ACE2 target protein’s motions (Figure 2C) the key functional sites of ACE2 that are recognized by the viral RBD are quite clearly and dramatically revealed by the MMD (Figure 3C). These include two sites on the N-terminal helices of ACE2 including two well documented additional sites identified at Q325 and K353 identified in previous studies of functional dynamics and evolution (Rajendran et al., 2022; Rajendran and Babbitt, 2022; Rynkiewicz et al., 2021).

In the fourth example, the DROIDS 5.0 KL divergence demonstrates a large allosteric effect of the eukaryotic specific InsP6 signaling molecule in triggering general amplification of dynamics across the whole of the Vibrio cholera RTX cysteine protease (Figure 2D), effectively activating the toxin only when it is present within host tissues thus preventing proteolytic destruction of the bacteria itself (Lupardus et al., 2008). The maxDemon 4.0 MMD analysis also captures this effect and additionally reveals the key cysteine and other possible sites that drive this change in dynamics (Figure 3D) .

To demonstrate the utility of MMD in a comparative evolutionary analysis of human vs. bacteria TBP (Figure 4), maxDemon 4.0 derived a neutral model distribution of MMD in dynamics for randomly selected pairs of differing amino acid sites on the human and bacterial orthologs (Figure 4A) with the tails indicating non-neutral evolution colored red for functionally conserved dynamics and green for adaptively altered dynamics. The two TBP ortholog structures are nearly identical (Figure 4B), and yet (Figure 4C) two regions of altered dynamics (i.e. high MMD) appear related amino acid replacements that have shifted the protein dynamics related to the TBP central hinge and one of the two loop binding regions highlighted earlier in Figure 3A. Most of the rest of the majority of the amino acid replacements (red bars) have occurred under the selective pressure to functionally conserve the TBP dynamics keeping the MMD low between the two orthologs.

The last ATOMDANCE method demonstrates the utility of a mixed effects model ANOVA combined with network community detection algorithms in ChoreoGraph 2.0 for identifying regions with coordinated or time resonant dynamics (Figure 5, Supplemental Figure 7) and for detecting similarities in dynamics driven by potential amino acid contacts (Supplemental Figure 8). Here we analyzed our fourth case example above comparing the unbound and InsP6-bound dynamics of the RTX cysteine protease domain. We demonstrate a profound loss of regions of resonant motions (i.e. communities of site resonance over time) that develops when allosteric activation occurs. Upon activation of the protease, the resonance shift from a tensed to a relaxed state mainly serves to remove most of the coordinated motions between amino acid sites. This was particularly exemplified by enhanced dynamics at the loop or flap regions at the top of the protease. This presumably would allow for more facile interactions with other protein substrates in the cell upon infection by the *V. cholera* pathogen. We have recently investigated protease flap dynamics in another protease system in HIV-1 (Rajendran et al., 2023). In this allosterically activated protease, we demonstrate another potentially valuable computational approach to analyzing allostery and other forms of protein logic gating in metabolic and regulatory pathways in the cell.

**Figure 4.**
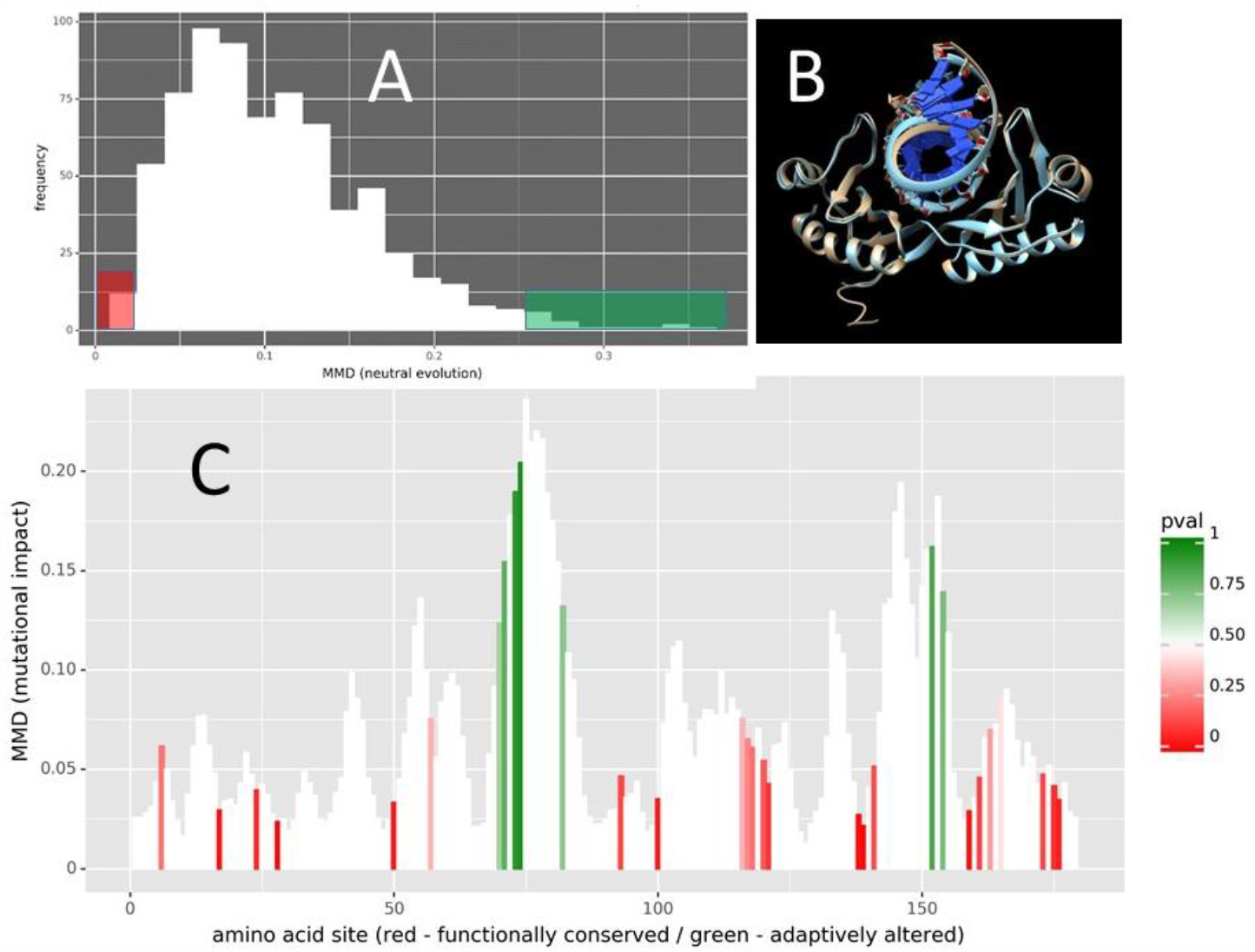
Site-wise unsigned maximum mean discrepancy MMD in local atom fluctuation and atom correlation comparing DNA-bound models of bacterial and human orthologs of TATA binding protein (PDB: 1qna and PDB 1cdw resp.). As a test of neutral evolution, the MMD between dynamics on randomly chosen differing amino acid sites between the orthologs is used to generate (A) an expected distribution of MMD for the effects of random amino acid replacement on molecular dynamics. The tails of the distribution are used to identify MMD values indicative of functionally conserved dynamics (red) or adaptively altered dynamics (green). (B) The superimposition of the two structures shows that the protein has maintained near perfect structural similarity since the divergence of common ancestor between bacteria and humans despite many amino acid replacements over time. (C) The MMD profile of the dynamic differences between orthologs is the background (in white) for the bootstrap analyses of MMD for the existing amino acid replacements (in color). Red indicates dynamic changes that are significantly smaller than expected under the neutral model (i.e. functionally conserved) while green indicates dynamic changes that are significantly larger (i.e. adaptively altered).

**Figure 5.**
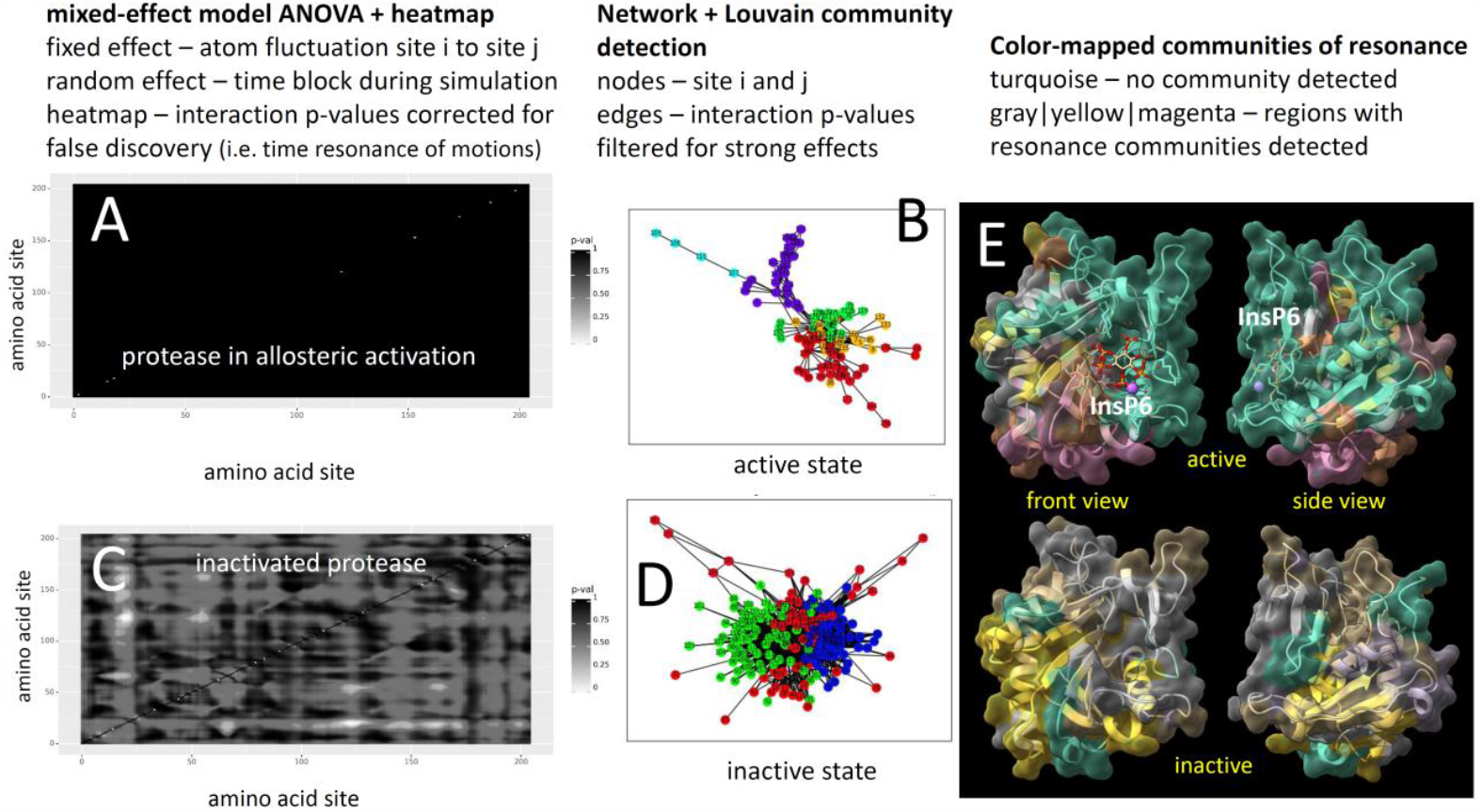
ChoreoGraph 2.0 - resonance analysis heat maps and community detection indicating regions of coordinated protein dynamics over time. The heat maps of the interaction term p-values for all pair-wise comparison of sites I to sites j on the (A) inactivated or unbound and (B) allosteric activated (i.e. InsP6 bound) Vibrio cholera toxin RTX cysteine protease domain (PDB: 3eeb) are shown. Multiple-test corrected interaction P-values for atom fluctuation across sites over time (i.e. resonance in motions) are derived from pair-wise mixed effects model ANOVAs where atom fluctuation is the dependent variable and sites i vs site j is the fixed effect and time samples are the random effect). Resonance patterns across sites are indicated by significant p-value (white). Regions of coordinated motion derived from Louvain community detection applied to graph network analysis are shown for (C) unbound and (D) InsP6-bound V. cholera protease. Resonance regions (i.e. communities of sites with significant time interactions) are similarly color mapped to the surface of the protein. Regions that fail to form resonance are colored light turquoise green. Note that upon binding (D) several very large resonance communities/regions disappear probably allowing the protease to more easily interact with host protein targets than when in the inactivated state within the bacteria. Connectivity and non-randomness of the resonance network is significantly higher in the inactivated state (connectivity t=-397.83, p<0.0001 | non-randomness t=-47.75, p<0.0001). Closeup view of the resonance heatmap for the inactivated protease (C) is given in Supplemental Figure 7.

## Discussion

Molecular dynamics simulation is a powerful tool for estimating physicochemical properties of systems in modern protein science. However, its utility has been limited by the lack of statistically sound methods that allow site-wise comparative functional and evolutionary analyses of protein dynamics. Unlike protein sequence and structural data, both static forms of data, capturing protein motion via molecular dynamics simulations creates a large component of variation that is induced by solvent-induced random thermal noise, subsequently creating a dataset in which non-random functional motions of proteins are obscured by this noise. We have described ATOMDANCE, a software suite for comparative protein dynamics that includes a powerful and interpretable kernel-based machine learning post-processor that allows users to mitigate the effects of noise and identify functional and evolutionary differences in molecular dynamics at individual sites on proteins.

While site-wise differences and divergence metrics can capture meaningful differences in protein function related to overall shifts in thermodynamics, they can have difficulty identifying the key binding sites without a large amount of time-consuming sampling via MD simulation. This is because dynamics at these sites, often characterized by non-random machine like motion, are potentially heavily masked by thermal noise in the MD simulation. The TATA binding protein used in our validations, is a perfect example of such a protein, as it utilizes two functional binding recognition loops, but nevertheless binds DNA so strongly so as to even bend the rigid DNA molecule and dampen atom fluctuation across nearly the whole of the TATA binding protein. While divergence metrics in DROIDS 5.0 capture this overall dampening at nearly all protein sites very well, our kernel learner in maxDemon 4.0 clearly identifies the functional binding sites themselves hidden within the thermal noise.

Finally, our application of mixed-effects model ANOVA combined with network graph analysis demonstrates a clear role of the coordination of motions of amino acid sites via time-dependent resonance in atom fluctuation in the allosteric control and logic-gating behavior of a well-studied pathogenic protease. We also note that this while our resonance analysis of this allosteric protease confirms the long hypothesized existence of ‘tensed’ and ‘relaxed’ protein states involved in allosteric regulation, it also represents a significant departure from past theory in that it demonstrates that these states are not always dependent upon interactions across separate protein domains that invoke conformational change(Guo and Zhou, 2016; Liu and Nussinov, 2016; Monod et al., 1965); however see (Cooper and Dryden, 1984). As we observe here, they can also occur across communities of coordinated sites acting within a single domain and in very rapid harmonic motion (i.e. < 1ns).

In conclusion, we have demonstrated the utility of ATOMDANCE for investigating a variety of functional aspects of protein systems including DNA-binding, drug/ligand induced shifts in dynamics, protein-protein interactions in infectious disease, as well as the functional evolutionary divergence/convergence of human-bacterial protein orthologs. ATOMDANCE offers traditional comparative metrics applied to molecular dynamics as well as a novel kernel-based approach to identifying specific key sites driving either binding interactions and/or protein activation. ATOMDANCE also offers analyses for characterizing regions across the protein where changes or shifts in dynamics over time are coordinated or time resonant across many sites. ATOMDANCE is entirely python-based with an easy to use graphical interface with seamless interaction with the open-source cpptraj MD analysis library and the modern UCSF ChimeraX molecular visualization software.

## Acknowledgements

We thank Dr. George M. Thurston for insightful feedback and creative suggestions for the early direction of this work.

## Supplemental File

video overview with dynamics of DNA-bound TATA binding protein and sorafenib drug-bound B-Raf kinase domain weighted in accordance with maximum mean discrepancy in atom fluctuation. https://people.rit.edu/gabsbi/img/videos/MMDmovie.mp4

