## Supplemental Figures for "ATOMDANCE: kernel-based denoising and allosteric resonance analysis for functional and evolutionary comparisons of protein dynamics"

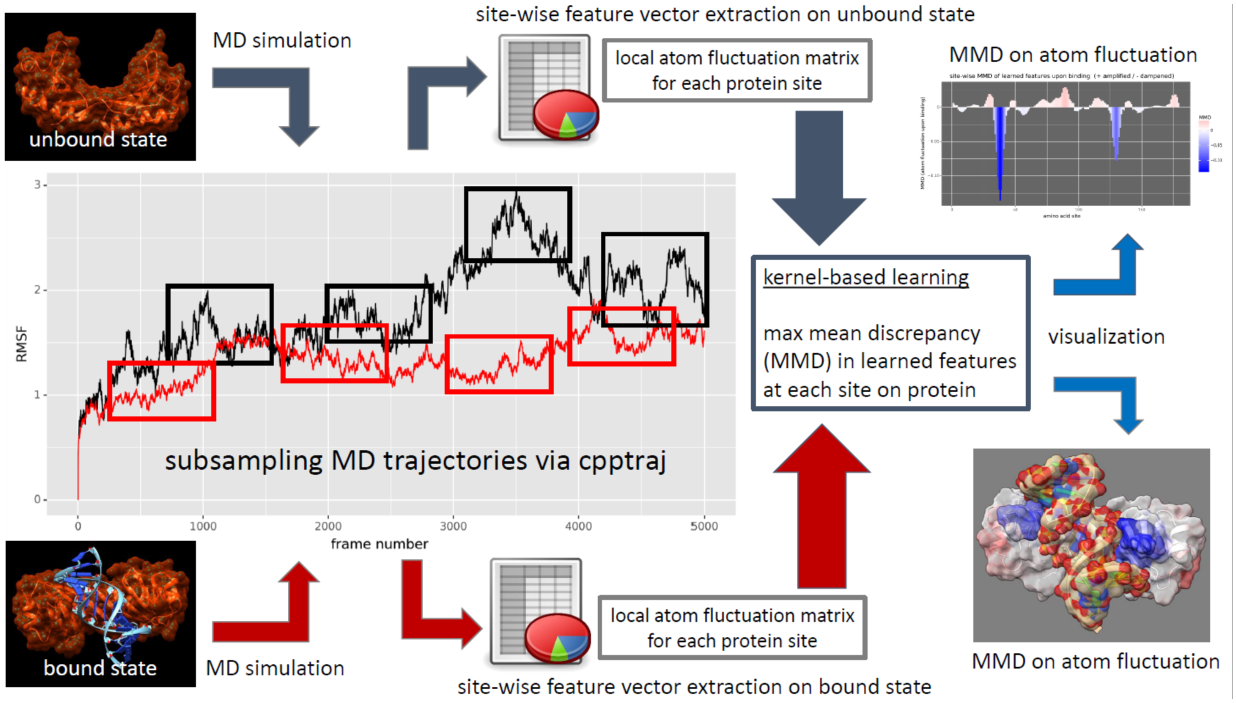


**Supplemental Figure 1 – Overview of the maxDemon 4.0 subroutine in the ATOMDANCE statistical machine learning post-processor for comparative protein dynamics.** First two molecular dynamics simulations representing the functional end states are conducted (e.g. drug/DNA/protein bound vs. unbound). The .pdb structure, .prmtop topology, and .nc trajectory files for both states are input to the software and the trajectories are repeated subsampled according to user specification using cpptraj. Site-wise local atom fluctuation matrices are constructed from the subsampling and used to train a Gaussian process kernel (radial basis function). At each site on the protein, the maximum mean discrepancy (MMD) in reproducing kernel Hilbert space is calculated, representing the distance between the learned features in the transformed data space that best captures the functional difference in protein dynamics at the given site. The MMD is signed negative if atom motion is dampened or positive if it is amplified. This MMD is visualized in a variety of plots and can be color-mapped to the .pdb structure file in UCSF ChimeraX (blue indicating regions of dampened motion due to binding interaction).

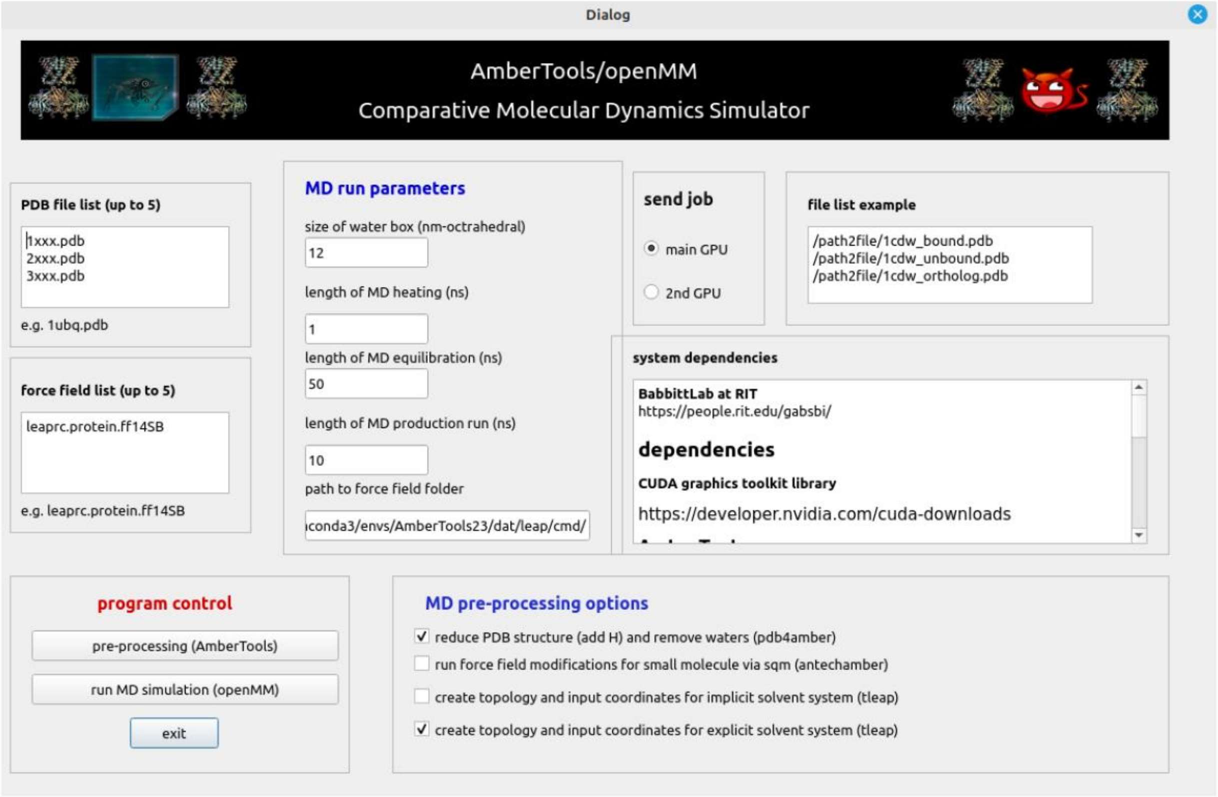


**Supplemental Figure 2 – Overview of the ATOMDANCE supplemental GUI for running MD simulations.** This program only requires a conda install for AmberTools and OpenMM. Users can batch run molecular dynamics (MD) simulations for up to 5 PDB structures by listing them in the window in the top left corner. The window below this should contain a list of all the necessary force fields (in AmberTools). Checkboxes for MD simulation pre-processing in AmberTools includes drying and reducing (i.e. adding hydrogens), calculating and defining force field modifications required by small molecule ligands (via antechamber and sqm = scaled quantum mechanical optimization), and preparing a charge neutralized solvated state by adding Na+ and Cl- ions and water molecules via tleap (in AmberTools).

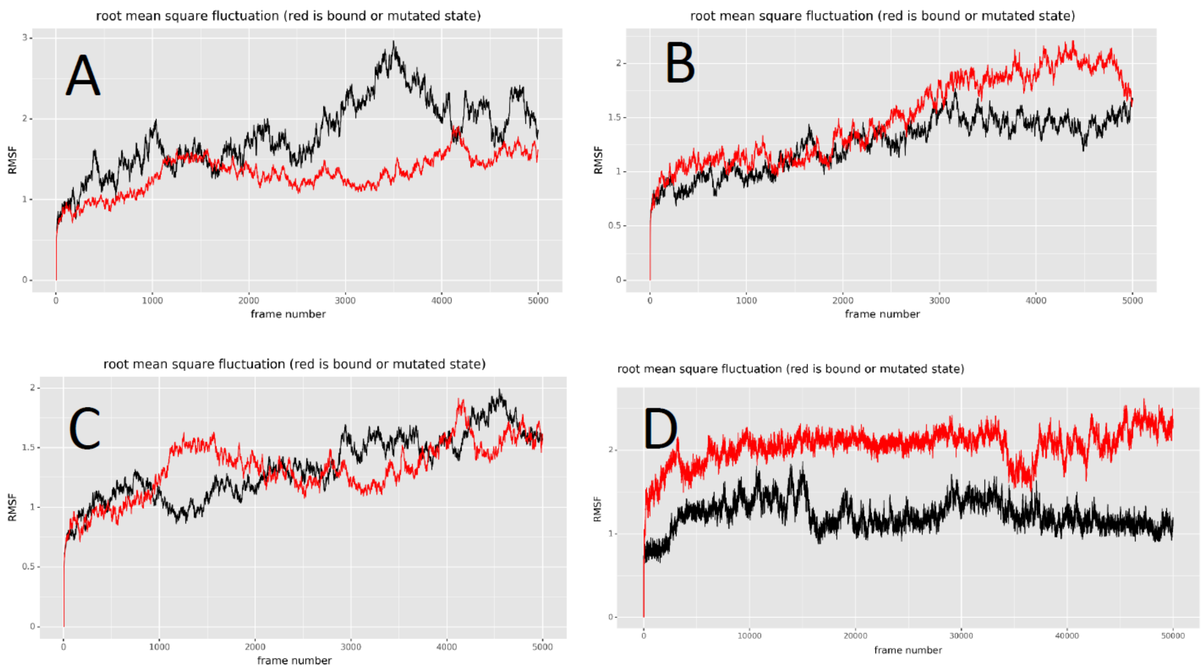


**Supplemental Figure 3 – Root mean square deviation over time (RMSF) plots for Figure 3 showing the 10ns production runs repeatedly sampled for the analyses.** The comparisons include (A) DNA-bound vs. unbound TATA binding protein (PDB: 1cdw), (B) sorafenib-bound vs. unbound B-Raf kinase domain (PDB: 1uwh), (C) SARS-CoV-2 viral bound vs. unbound angiotensin-converting enzyme 2 (ACE2) protein (PDB: 6m17), and (D) the allosteric activated (i.e. InsP6 bound) vs inactivated (i.e. unbound) Vibrio cholera toxin RTX cysteine protease domain (PDB: 3eeb). The RMSF for the molecular dynamics trajectory over time for unbound state of the target proteins are shown in black and the bound state is shown in red.

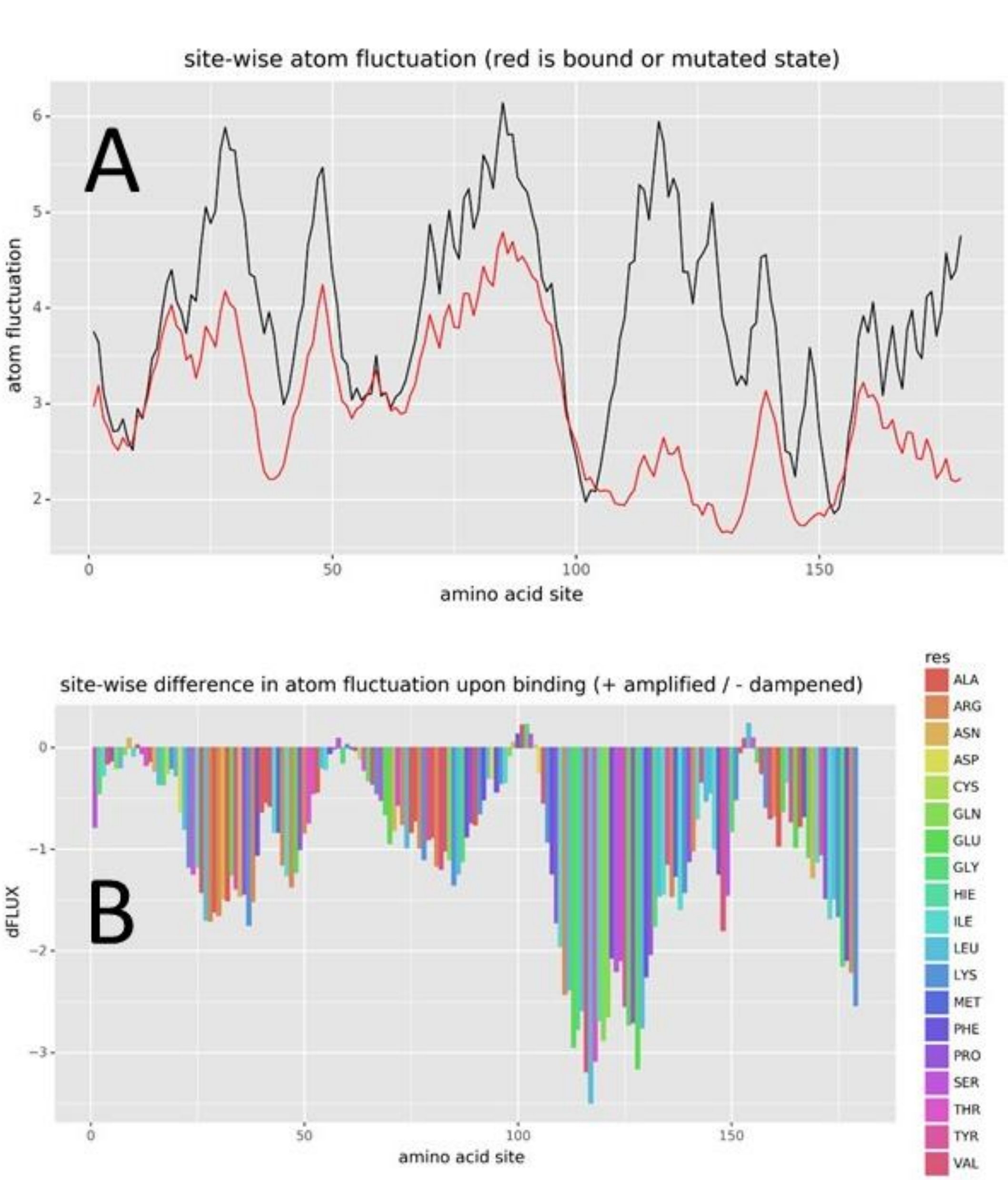


**Supplemental Figure 4 – Alternative plots generated by ATOMDANCE** indicating (A) site-wise average atom fluctuation profiles and (B) site-wise average difference in DNA-bound versus unbound TATA binding protein (PDB: 1cdw).

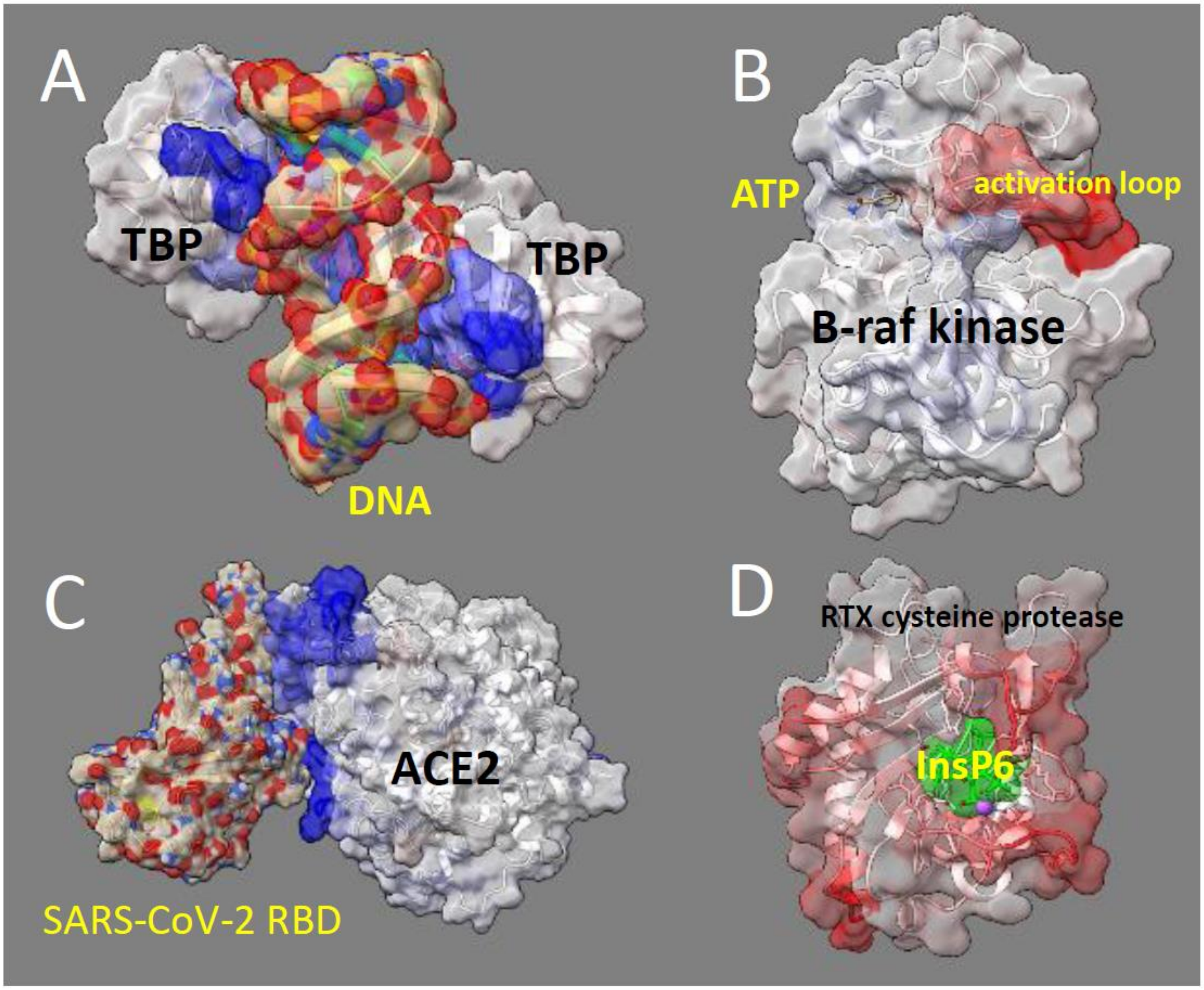


**Supplemental Figure 5 – Close-up views of color-mapped structures from Figure 3.** Note red indicates the maximum mean discrepancy (MMD) between learned features where atom motions are amplified in the ligand-bound state and conversely, blue indicates where these atom motions are dampened. The comparisons include (A) DNA-bound vs. unbound TATA binding protein (PDB: 1cdw), (B) sorafenib-bound vs. unbound B-Raf kinase domain (PDB: 1uwh), (C) SARS-CoV-2 viral bound vs. unbound angiotensin-converting enzyme 2 (ACE2) protein (PDB: 6m17), and (D) the allosteric activated (i.e. InsP6 bound) vs inactivated (i.e. unbound) Vibrio cholera toxin RTX cysteine protease domain (PDB: 3eeb).

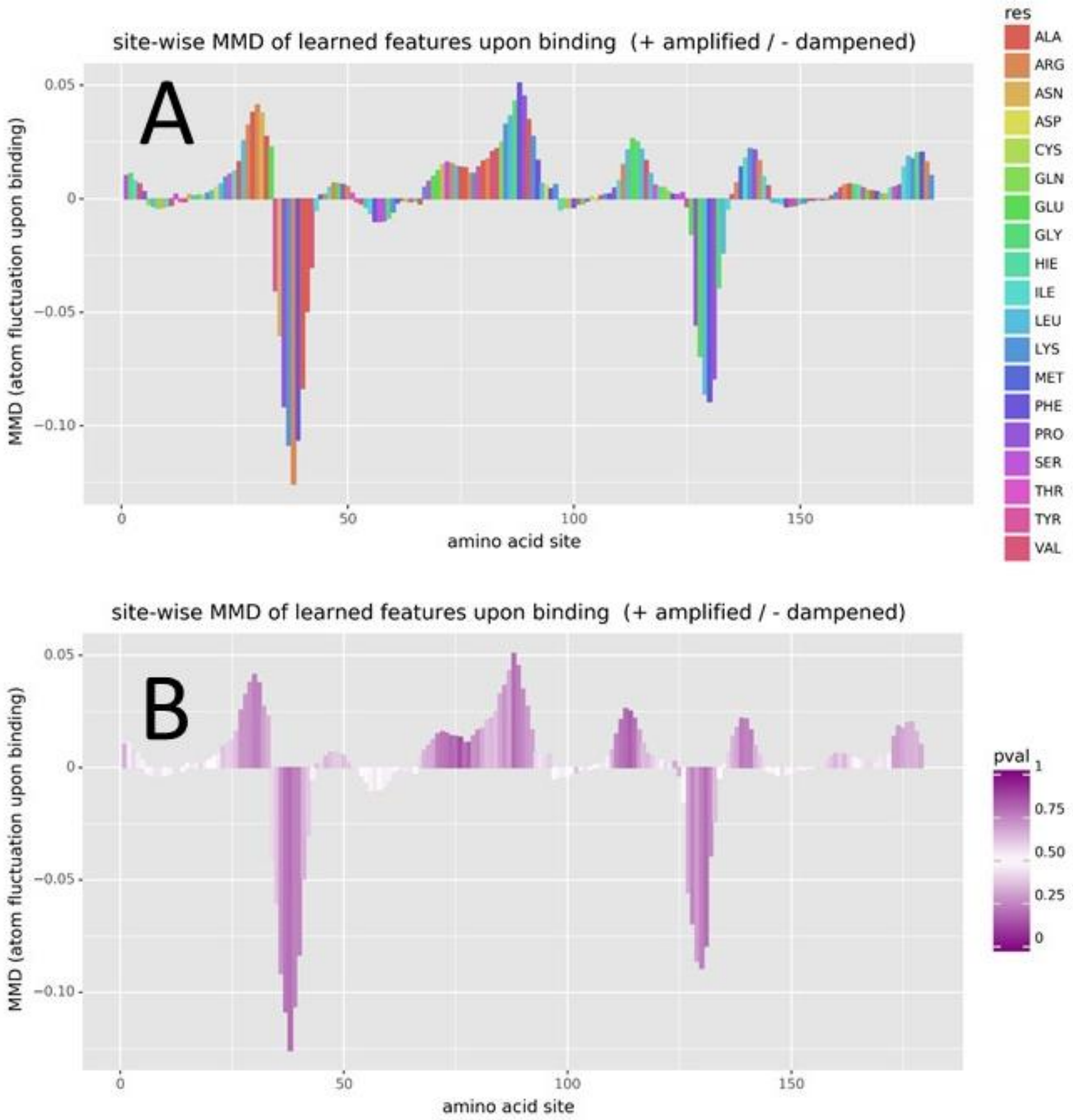


**Supplemental Figure 6 – Alternative plots generated by ATOMDANCE** indicating site-wise max mean discrepancy (MMD) in learned features trained on local atom fluctuation showing alternative color plotting for (A) amino acid type and (B) bootstrapped empirical p values.

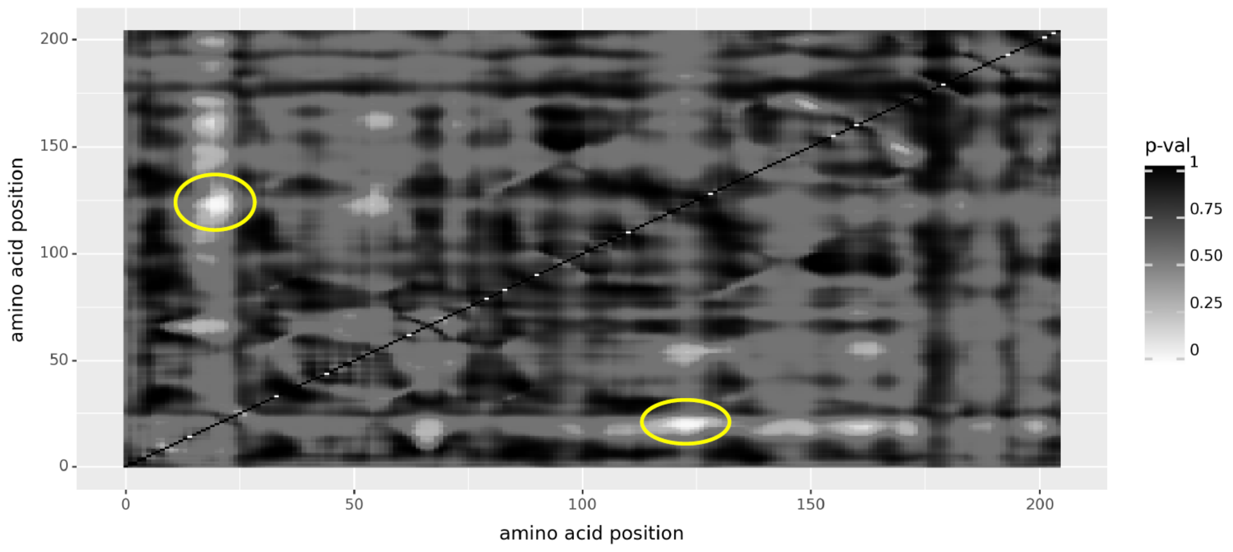


**Supplemental Figure 7 – Close-up of the heatmap of significant site resonance during allosteric inactivation of RTX cysteine protease in the absence of InsP6 (shown in Figure 5C).** Significant resonance between sites is indicated by interaction p-values (corrected for false discovery rate) derived from a mixed-effects model ANOVA where atom fluctuations at sites I and j and the fixed effect in the model and the time during the molecular dynamics simulation is the random effect. Note: allosteric activation by InsP6 removes all signatures of significant resonance.

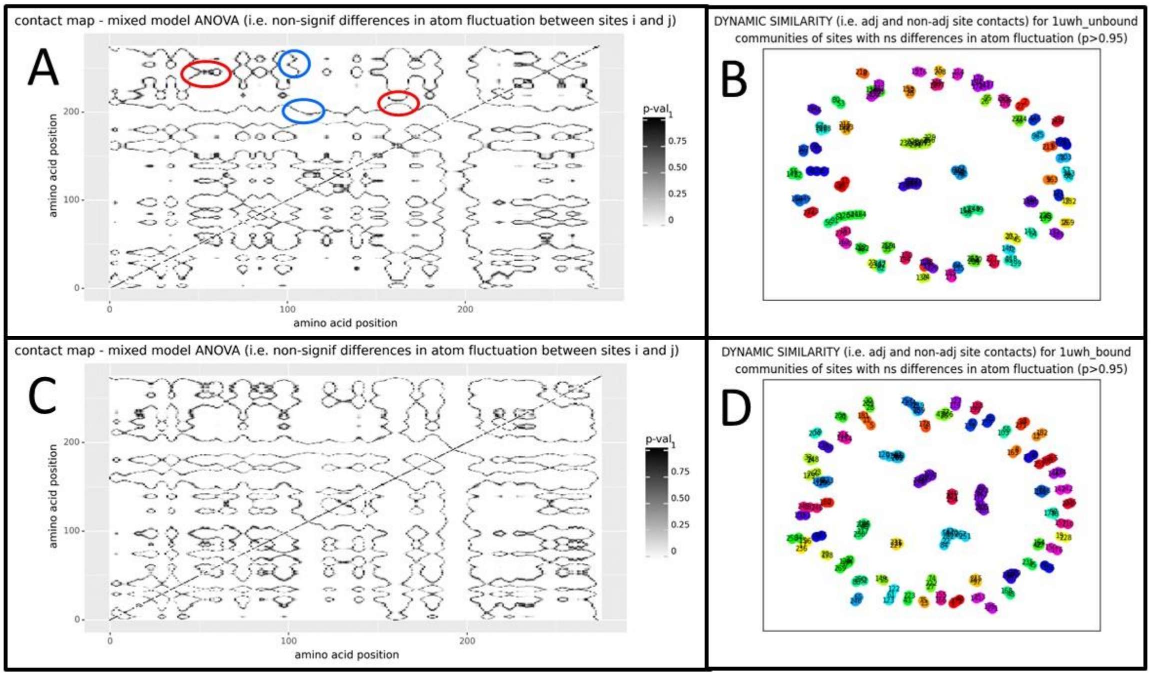


**Supplemental Figure 8 – Potential contact analysis heat maps and community detection (ChoreoGraph 2.0) indicating regions of overall similarity of protein dynamics** (regardless of time). The heat maps of the fixed effect term p-values for all pair-wise comparison of sites I to sites j on the (A) unbound and (C) drug sorafenib-bound BRAF kinase protein (PDB: 1uwh) are shown. Non-significant p-values (black) indicate overall similarity of atom fluctuation between adjacent or non-adjacent sites. Some examples of patterns of similarity caused by adjacent sites (circle blue) and non-adjacent contacts (circled red) are shown in (A). The latter are identified where the non-significant p-value trace on the map doubles back and connects distant sites. Regions of overall similar motion derived from Louvain community detection applied to graph network analysis are shown for (B) unbound and (D) sorafenib-bound BRAF kinase. Similar regions (i.e. communities of sites with non-significant differences in overall atom fluctuation) tend to form separate communities, often only pairs or triplets of sites under the community detection algorithm.

Supplemental File – video overview with dynamics of DNA-bound TATA binding protein and sorafenib drug-bound B-Raf kinase domain weighted in accordance with maximum mean discrepancy in atom fluctuation. https://people.rit.edu/gabsbi/img/videos/MMDmovie.mp4
