## Supplemental Methods for "ATOMDANCE: kernel-based denoising and allosteric resonance analysis for functional and evolutionary comparisons of protein dynamics"

**PDB structure and model preparation**

Protein structures of ubiquitin, TATA binding protein (TBP) (PDB: 1cdw)(Nikolov et al., 1996), BRAF kinase bound to the drug sorafenib (PDB: 1uwh)(Wan et al., 2004), SARS-CoV-2 receptor binding domain bound to angiotensin converting enzyme 2 (ACE2)(PDB:6m17)(Yan et al., 2020), and the allosteric RTX cysteine protease domain of the Vibrio cholera toxin (PDB:3eeb)(Lupardus et al., 2008) were obtained from the Protein Data Bank (PDB). After downloading the structures from the PDB database, any crystallographic reflections, ions, and other solvents used in the crystallization process were removed. Any missing loop structures in the protein structures were inferred using the MODELLER homology modelling server in UCSF Chimera. pdb4amber (AmberTools20) was employed to add hydrogen atoms (i.e. reduce the structure) and remove crystallographic waters.

**Molecular dynamic simulation protocols**

For each molecular dynamic comparison (monomer vs. dimer, wildtype vs. mutant protease; protease bound and unbound to drug), accelerated molecular dynamic (MD) simulations were performed (Case et al., 2005b). MD simulation protocol was followed as previously described, with slight modifications(Babbitt et al., 2022a, 2022b, 2022c, 2020, 2020, 2018; Rajendran et al., 2022; Rajendran and Babbitt, 2022). In brief, for each MD comparison, large replicate sets of accelerated MD simulation were prepared and then conducted using the particle mesh Ewald method implemented on A100 and V100 NVIDIA graphical processor units by pmemd.cuda running Amber20 (Case et al., 2005a; Darden et al., 1993; Ewald, 1921; Pierce et al., 2012; Salomon-Ferrer et al., 2013) and/or OpenMM (Eastman et al., 2017). The MD simulations were done on a high performance computing workstation mounting dual Nvidia 2080Ti graphics processor units. All comparative MD analysis via our ATOMDANCE was based upon 50 randomly resampled windows collected on of 10ns of accelerated MD in each comparative state, e.g., monomer vs. dimer, wildtype vs. mutant, protease bound to drug vs. protease unbound to drug). Explicitly solvated protein systems were first prepared using teLeap (AmberTools 20), using ff14SSB protein force field, in conjunction with modified GAFF2 small molecule force field (Maier et al., 2015; Wang et al., 2004). Solvation was generated using the Tip3p water model in a 12nm octahedral water box. Charge neutralization was performed using Na+ and Cl- ions using the AmberTools22 teLeap program. Force field modifications for the small molecule ligands were generated using scaled quantum mechanical optimization via the sqm version 17 program in antechamber/AmberTools22 (Walker et al., 2008). For each MD comparison, an energy minimization was first performed, then heated to 300K for 300 pico seconds, followed by 10 ns of equilibration, and then finally a replicate set of 100 MD production run was created for each comparative state. Each MD production run was simulation for 1 ns of time. All simulations were regulated using the Anderson thermostat at 300k and 1atm (Andersen, 1980). Root mean square atom fluctuations were conducted in CPPTRAJ using the atomicfluct command (Roe and Cheatham, 2013). All molecular color-mapping of our results were conducted in UCSF ChimeraX (Goddard et al., 2018; Pettersen et al., 2021). Any x-ray crystal protein structures requiring missing loop refinement were corrected using MODELLER prior to preparation for molecular dynamics simulations (Sali and Blundell, 1993).

ATOMDANCE.py is a PyQt5 GUI designed for post-processing comparative molecular dynamics and delivering information about important protein site differences between the dynamics of proteins in two different functional states. It also can be used to investigate potential site-wise evolutionary changes in protein dynamics and to investigate where sites share coordinated dynamics states as well. After randomly subsampling the atom trajectory files and calculating amino acid site atom fluctuations and atom correlations using the atomicfluct and atomiccorr functions from the cpptraj library, ATOMDANCE.py runs 4 types of analyses listed below.

DROIDS 5.0 – protein site-wise divergence metrics for pair-wise comparison of protein backbone atom fluctuations across functional protein states (e.g. bound vs. unbound or wildtype vs. mutant)

This option is an acronym for Detecting Relative Outlier Impacts in Dynamics Simulations and calculates both the average differences and KL divergences in the atom fluctuation at every protein site. Fluctuations are averaged over the four protein backbone atoms for each amino acid (i.e. N, Cα, C, and O). Significant differences in dynamics of the two protein states are determined by a two sample Kolmogorov-Smirnov test corrected for the number of sites in the protein corrected for the false discovery rate (i.e. Benjamini-Hochberg method) caused by the total number of sites on the protein. This method is described and published previously in DROIDS v1.0-4.0 (Babbitt et al., 2020, 2018). The only difference in v5.0 is that the subsampling is taken from random window positions along a single long MD production run, rather than multiple short MD production runs.

maxDemon 4.0 – protein site-wise kernel learning for pair-wise comparison of protein backbone atom fluctuations and/or atom correlations across functional protein states (e.g. bound vs. unbound or wildtype vs. mutant)

This analysis option uses site-wise training of Gaussian processes machine learners with tuned radial basis kernel functions in order to specify a maximum mean discrepancy (MMD) in reproducing kernel Hilbert space (RKHS) that describes the distance in learned features between the two protein dynamic states at all given sites on the protein.

Thus the kernel function describing the mapping of the data points $x_i$ and $x_j$ being compared is

$$k(x_i , x_j ) = exp\left(\frac{\|x_i - x_j\|^2}{2\sigma^2}\right)$$

And the empirical estimations of MMD, or distance between feature means is given by

$$MMD^2(X,Y) = \frac{1}{m(m-1)}\sum_{i}\sum_{j\neq i} k(x_i, x_j) - 2\frac{1}{m(m-1)}\sum_{i}\sum_{j} k(x_i, y_i) + \frac{1}{m(m-1)}\sum_{i}\sum_{j\neq i} k(y_i, y_j)$$

where x's are the data points we have and y's are generated examples evaluated on the kernel.

The learners in ATOMDANCE can be trained using a local atom fluctuation feature vector comprised of fluctuations from sites -2, -1, 0, 1, 2 positions on the protein chain relative to the site being analyzed. The observed site-wise MMD values are further subjected to hypothesis testing using a bootstrap derived empirical p-value whereby the observed MMD values between the functional dynamic states at any given site are compared to 500 bootstrapped MMD values for that site when derived from resampling the dynamics in the same dynamic state. A graphical overview of this analysis is shown in Figure 1. A key concept here when comparing this output to the site-wise KL divergence metrics generated by DROIDS 5.0 is that because the learner cannot optimize on random differences in atom fluctuation caused by thermal noise it acts as a noise filter, thus eliminating motion dampening that is not directly due to non-random differences in atom fluctuation between the sites being compared (i.e. functional aspects of molecular interactions directly involved in the binding interaction)

maxDemon 4.0 – protein site-wise kernel learning for detecting natural selection acting upon protein dynamics via neutral modeling of amino acid replacements (e.g. human vs. another species ortholog or another human paralog)

This analysis option is only appropriate when comparing two homologous proteins in two states of molecular evolution, whereby mutations have accrued over time and the user would like to determine whether the dynamics at a given site of amino acid replacement has likely been functionally conserved, evolved neutrally or evolved adaptively (i.e. under purifying, neutral or adaptive evolution). Comparisons of dynamics between the same protein in two different species (i.e. orthologs) or two related proteins in the same species (i.e. paralogs) are both enabled through this method of analysis. In this case, the MMD in dynamics between each site of amino acid replacement between the homologous proteins is compared to a model of neutral evolution represented by a distribution of MMD taken from the dynamics of 500 random pairs of dissimilar amino acids at different sites on the homologous proteins. If the MMD for an observed amino acid replacement is in the lower or upper extremities of the distribution of neutral MMD (two-tailed level of significance = 0.05), then natural selection acting upon the dynamics can likely be inferred.

ChoreoGraph 2.0 – identification of protein site communities with coordinated dynamic states mixed-effects model ANOVA and Louvain community detection

This analysis option examines the reference and query dynamic state simulations of the proteins compared above and produces (A) site-wise heatmaps and community level graph networks identifying groups of amino acid sites where atom fluctuation values are resonating over time in a coordinated fashion and (B) site-wise heatmaps and community level graph networks identifying groups of amino acid sites where overall atom fluctuation values are not significantly different from each other (i.e. potentially in contact). In each dynamic state simulation every site *i* on the protein is compared to every site *j* using a mixed effects model ANOVA where atom fluctuation represents a fixed effect (α) in the model and a time sample represents a random effect (β) in the model. Thus the general linear model becomes

$$Y_{st} = \mu + \alpha_s + \beta_t + \alpha\beta_{st} + \varepsilon_{st}$$

where *s* represents the site class (*i* or *j*) and *t* represents the random time sampling group collected by the cpptraj program (see subsampling step in Figure 1).

For the resonance analysis, the p-value of interaction between atom fluctuation levels between site *i* and site *j* and the time subsamples in the MD simulation (i.e. $\alpha\beta_{st}$) indicates the significance of an interaction of atom fluctuation between the two different sites over time (i.e. a coordinated physical resonance in motion). The p-values corrected for false discovery rate via Benjamini Hochberg method are shown as a heatmap representing significant areas of site resonance on the protein. In the second step of the analysis, intended to define communities of resonating regions of protein dynamics, the uncorrected interaction p-values for all site *i* to site *j* comparisons are represented by a graph network where ($k_n$), the degree of node n is

$$k_n = A_{nm} \sum m$$

where *n* and *m* are the interaction p values for sites *i* and *j,* and $A_{nm}$ is the adjacency matrix connecting nodes *n* and *m*. The Louvain community detection algorithm (Blondel et al., 2008) iterates a two step process of modularity optimization followed by community aggregation until community identities of all nodes are stable.  It is implemented in our code by the python package networkx (Hagberg et al., n.d.).

For the analysis of potential adjacent and non-adjacent contacts, the same procedure as used above is repeated with the exception that the p-values no longer represent the interaction terms of the ANOVA, but now represent the fixed effect of the model. So in this case the resulting heatmap and network communities represent sites whose overall atom fluctuations are NOT significantly different from each other.  In general, these networks are highly fragmented (i.e. often pairs of sites) and a trace of the non-significant p-values in the heatmap can easily be used to assess whether sites *i* and *j* are adjacent or non-adjacent to each other.

ATOMDANCE is available at GitHub/GitHub page

https://github.com/gbabbitt/ATOMDANCE-comparative-protein-dynamics

https://gbabbitt.github.io/ATOMDANCE-comparative-protein-dynamics/

Examples presented in this manuscript were generated from structure, topology, and trajectory files deposited here

https://zenodo.org/record/7679282#.Y_wIK9LMJ9A

DOI 10.5281/zenodo.7679282

makeMovie.py

A supplemental python GUI program for making molecular dynamics movies that are weighted by the normalized MMD in atom fluctuation between two functional states. The program first creates a multi-frame PDB file representing the true dynamics of the protein system, then it creates a multi-frame PDB file where the noise in the trajectories is dampened or amplified according to MMD. This creates a purely visual effect in a color-mapped movie of protein motion that demonstrates what the MMD filter captures. We have demonstrated this in examples of both dampening of atom motion during TATA binding protein interaction with DNA and with amplification of motion in the activation loop of BRAF kinase during cancer drug binding. https://people.rit.edu/gabsbi/img/videos/MMDmovie.mp4

MDgui.py

We also provide a full python GUI for running MD simulations using open source AmberTools and openMM. The user can generate MD trajectory and topology files using any software they prefer. Other options include Amber (licensed), NAMD/QwikMD (free), CHARMM (licensed), or openMM (free). The cpptraj software available on GitHub or in AmberTools can be used to convert common trajectory file formats to the binary format (.nc) used by ATOMDANCE. We also offer a useful python+perl-based GUI for licensed versions of Amber available here. https://gbabbitt.github.io/amberMDgui/

Note on the naming of things:
DROIDS – acronym for Detecting Relative Outlier Impacts in Dynamics Simulations
maxDemon – abbreviated from Maxwell's Demon, a 19$^{th}$ century thought experiment connecting the concepts of information and entropy in thermodynamics involving a mythical demon watching/assessing the motion of every atom in a system.
ChoreoGraph – evokes a notion of when motions of atoms at amino acids site 'move together' in a coordinated manner, in much the same way dancers may move together in choreography.
ATOMDANCE – an homage to a song composition by Icelandic singer Bjork Guomundsdottir from her 2015 album Vulnicura (One Little Indian Records)

Supplemental File – video overview with dynamics of DNA-bound TATA binding protein and sorafenib drug-bound B-Raf kinase domain weighted in accordance with maximum mean discrepancy in atom fluctuation. https://people.rit.edu/gabsbi/img/videos/MMDmovie.mp4
